# Genetic variation in the social environment affects behavioral phenotypes of oxytocin receptor mutants in zebrafish

**DOI:** 10.1101/2020.03.23.002873

**Authors:** D. Ribeiro, A.R. Nunes, M.C. Teles, S. Anbalagan, J. Blechman, G. Levkowitz, R.F. Oliveira

## Abstract

Oxytocin-like peptides have been implicated in the regulation of a wide range of social behaviors across taxa. On the other hand, the social environment, which is composed of conspecifics genotypes, is also known to influence the development of social behavior, creating the possibility for indirect genetic effects. Here we used a knockout line for the oxytocin receptor in zebrafish to investigate how the genotypic composition of the social environment (E_s_) interacts with the oxytocin genotype (G) of the focal individual in the regulation of its social behavior. For this purpose, we have raised wild-type or knock-out zebrafish in either wild-type or knock-out shoals and tested different components of social behavior in adults. GxE_s_ effects were detected in some behaviors, highlighting the need to control for GxE_s_ effects when interpreting results of experiments using genetically modified animals, since the social environment can either rescue or promote phenotypes associated with specific genes.

## Introduction

Social genetic effects (aka indirect genetic effects) occur when the phenotype of an organism is influenced by the genotypes of conspecifics. Previous work has highlighted the major potential evolutionary consequences of social genetic effects(1,2), with evidence for such effects to be present both in interactions between related (e.g. mothers and offspring(3,4)) and unrelated individuals (e.g. sexual displays(5), aggression (6–8)). More recently the importance of social genetic effects for health and disease has also been recognized (9), which may explain the pervasiveness of the social environment as a mortality risk in humans (10,11). Interestingly, the potential consequences of social genetic effects for the interpretation of research results using genetically modified organisms (GMO) has been greatly neglected. GMOs have been widely used in behavioral neuroscience to investigate the causal role of candidate genes and behavioral phenotypes. Typically Knock-in and Knock-out transgenics and mutants have been used to causally link the gain or loss of behavioral function to a specific gene (12). In recent years the development of genome editing techniques, such as CRISPR-Cas9-and TALEN-induced mutations, have increased the interest in this approach and opened the door to studying the genetic basis of behavior in non-model organism (13). However, most studies using GMO in behavioral neuroscience have ignored the potential contribution of the environment to the behavioral phenotype studied. This is because it has been assumed that if the genetic background of these mutants is identical and their environment has been kept constant, any phenotypic differences must come from the genetic manipulation. However, when GMOs are incrossed or visually screened at a very young age (e.g. using reporter genes, GFP) and thereafter raised and housed together until used in experiments, changes in their behavior might be affected by divergent social environments experienced by these mutants. In other words, modified behavior might be as a result of growing with their peer mutants, rather than the canonical social environment provided by wild type conspecifics. Such problem is particularly relevant when studying social behavior. For example, mutant *Drosophila* males that lack the fruitless gene (*fru^M^*) do not express courtship behavior when held in isolation, but acquire the potential to court females when grouped with other flies (14), suggesting the occurrence of a combined gene (G) and social environment (E_s_) effect (aka, GxE_s_). Thus, given the rising interest in the study of social behavior in model organisms from worms to higher vertebrates, an assessment of potential gene by social environment effects on the behavioral phenotype of interest in GMOs used in social neuroscience is crucial).

Despite the wide variety of species-specific social behaviors, a wealth of evidence has implicated the paralog nonapeptides vasopressin (VP) and oxytocin (OXT) and their receptors in the regulation of different aspects of social behavior across vertebrates(15,16), suggesting a genetic toolkit role for these nonapeptides in social behavior. Nonapeptides are an ancestral neuropeptide family found both in vertebrates and invertebrates, that derived from a VP-like peptide, and that evolved along two parallel clades of VP- and OXT-like peptides derived from the duplication of the VP gene in early jawed fish (ca. 500 Mya). Both peptides have been implicated in the regulation of behavior and physiology across different taxa, with VP being more involved in aggression and agonistic behaviors and OXT-like peptides consistently acting in affiliative behaviors and species-specific social behaviors across diverse taxa (i.e. sexual behavior, social interactions) (17,18). Despite this wealth of evidence on the direct genetic effects of OXT on social behavior, social genetic effects of OXT genotypes have never been studied.

In this study we aimed to provide a proof of principle for GxE_s_ effects in behavioral phenotypes observed in GMO by assessing the occurrence of such effects in a knockout line for the OXT receptor in zebrafish, a commonly used model species in behavioral neuroscience (19). For this purpose, we studied the GxE_s_ interaction in the effects of the OXT gene (*oxtr*) in different aspects of social behavior, namely: social preference (i.e. social approach); social habituation; social recognition; and shoaling behavior. For this purpose, we have raised individual zebrafish of the WT (*oxtr^(+/+)^*) or knock-out genotype (*oxtr^(-/-)^*) in different social environments (i.e. *oxtr^(+/+)^* shoal or *oxtr^(-/-)^* shoal; Figure 1A).

**Figure 1.**
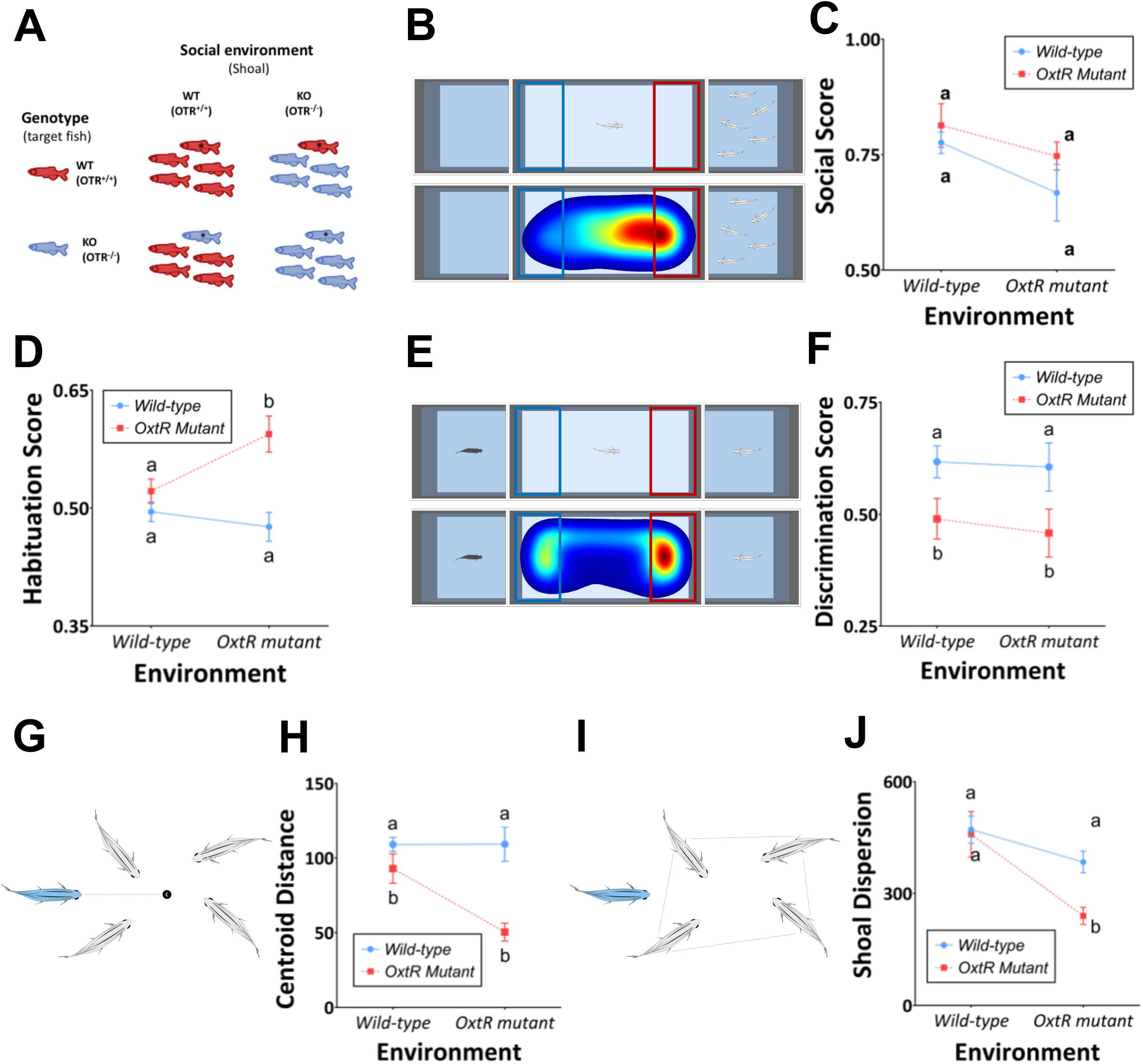
The contribution of the social environment (E_s_) and genotype (G) to the expression of behavioral phenotypes in zebrafish was assessed by raising oxytocin receptor mutant fish and wild types (focal fish marked with *) in shoals of either mutants or wild types (A). Social preference, measured by the time spend near a shoal in a choice test (B), showed no effect of either G or E_s_ (C). Social habituation, which consisted on a consecutive social preference test exhibited a GxE_s_ effect (D). Social recognition, measured as the discrimination between a novel and a familiar conspecific (E), shows a pure G effect (F). Social integration, measured as distance to the centroid of the shoal (G), showed a GxE_s_ effect (H). Social influence, measured by the cohesion of the remaining shoal members (I), also showed a GxE_s_ effect (J). Data is presented as mean ± standard error of the mean (SEM). Sample sizes are 9 for heterogeneous groups (i.e. focal individual with different genotype from the remaining individuals in the shoal; mutant focal in WT shoals and WT focal in mutant shoals) and 15 for homogeneous groups (i.e. focal individual with the same genotype of the remaining individuals in the shoal; mutant focal in mutant shoals and WT focal in WT shoals). Different letters indicate significant differences (p<0.05) between treatments as assessed by Two-Way ANOVA followed by post-hoc tests (see Table 1).

## Results and Discussion

Adult zebrafish, like many other social animals, express a tendency to approach and interact with conspecifics (social preference, Figure 1B) (20). Here we show that there was no significant effect of either genotype, environment or GxE_s_ interaction on social preference (Table 1; Figure 1C). However, when fish were presented for a second time to a shoal to measure social habituation (i.e. expected reduction in social preference), we found a GxE_s_ interaction, where *oxtr^(-/-)^* individuals raised in *oxtr^(-/-)^* shoals express enhanced social habituation (F_1,44_ = 5.642, p = 0.022; Figure 1D). In contrast, when we tested social recognition, which is a form of social memory needed for individuality in social interactions (i.e. differential expression of social behavior depending on identity of interacting individual), that is known to be modulated by oxytocin both in mammals and zebrafish (21,22), we observed that *oxtr^(-/-)^* individuals exhibit a deficit in acquisition and retention of social recognition irrespective of the social environment (*oxtr^(-/-)^* or *oxtr^(+/+)^*) in which they were raised (F_1,44_ = 7.600, p = 0.008; Figure 1F).

**Table 1.** Two-way ANOVA for ‘genotype’, ‘environment’ and their interaction (‘genotype’ x ‘environment’) was analysed for each behavioural essay. * indicates p < 0.05, ** indicates p < 0.01, *** indicates p < 0.001.

| **Social Preference** |  |  |  |  |
| --- | --- | --- | --- | --- |
|  | d.f. | Mean Squares | F | Significance |
| **Genotype (G)** | 1 | 0.023 | 1.731 | 0.195 |
| **Environment (E)** | 1 | 0.050 | 3.788 | 0.058 |
| **G x E** | 1 | 0.001 | 0.049 | 0.825 |
| **Error** | 44 | 0.013 |  |  |
| **Habituation** |  |  |  |  |
|  | d.f. | Mean Squares | F | Significance |
| **G** | 1 | 0.058 | 13.927 | 0.001 ** |
| **E** | 1 | 0.008 | 1.936 | 0.171 |
| **G x E** | 1 | 0.024 | 5.642 | 0.022 * |
| **Error** | 44 | 0.004 |  |  |
| **Social Recognition** |  |  |  |  |
|  | d.f. | Mean Squares | F | Significance |
| **G** | 1 | 0.213 | 7.600 | 0.008 ** |
| **E** | 1 | 0.005 | 0.189 | 0.666 |
| **G x E** | 1 | 0.001 | 0.041 | 0.841 |
| **Error** | 44 | 0.028 |  |  |
| **Social Competence** |  |  |  |  |
|  | d.f. | Mean Squares | F | Significance |
| **G** | 1 | 39.486 | 24.370 | <0.001 *** |
| **E** | 1 | 12.565 | 7.755 | 0.008 ** |
| **G x E** | 1 | 12.811 | 7.907 | 0.007 ** |
| **Error** | 44 | 1.620 |  |  |
| **Social Group Dispersion** |  |  |  |  |
|  | d.f. | Mean Squares | F | Significance |
| **G** | 1 | 174.366 | 4.309 | 0.044 * |
| **E** | 1 | 657.221 | 16.240 | <0.001 *** |
| **G x E** | 1 | 122.980 | 3.039 | 0.088 |
| **Error** | 44 | 40.469 |  |  |

Given that social behavior of zebrafish mainly occurs in the context of shoaling we have also investigated two shoaling behavior parameters: (1) social integration – how well the focal individual integrates in the shoal, as measured by its average distance to the centroid of the shoal (Figure 1G,H); and (2) social influence – how the focal individual affects the shoaling behavior of the remaining shoal members, by measuring the shoal dispersion as defined by the perimeter of the other shoal members (Figure 1I,J). A GxEs interaction was found for social integration: *oxtr^(-/-)^* individuals raised in *oxtr^(-/-)^* shoals exhibit a significantly lower social integration than *oxtr^(-/-)^* individuals raised in *oxtr^(+/+)^* shoals; in contrast, *oxtr^(+/+)^* individuals exhibit high levels of social integration irrespective of the shoal type in which they were raised (Table 1; Figure 1H). Finally, the presence of a single WT *oxtr^(+/+)^* individual in a *oxtr^(-/-)^* shoal was enough to increase its dispersion, whereas the presence of a single *oxtr^(-/-)^* individual in a *oxtr^(+/+)^* shoal did not affect its dispersion (Table 1; Figure 1J).

In summary, we show that distinct components of social behavior are differentially affected by the genetic composition of the social environment versus the *oxtr* genotype of the focal individual. Social recognition exhibited a pure effect of the genotype. However, clear GxE_s_ interactions were observed in the cases of social habituation and social integration, and social influence had a major contribution of the social environment. Thus, we demonstrated that genetic variation in the social environment interacts with individual genotype during the developmental acquisition of social behavior. In other words, the social environment can revert phenotypes associated with specific genes. These results are in line with reported interactions between the social environment and oxytocin receptor genotype in the determination of social behavior phenotypes in human populations (23–25). Our results suggest that more caution is needed in the interpretation of studies using transgenic or mutant individuals that are raised in cohorts of the same genotype, and that some phenotypes observed in transgenic or mutant lines may in fact result from GxES interactions.

## Materials and Methods

### Zebrafish lines and maintenance

Zebrafish were raised and bred according to standard protocols and all experimental procedures were approved by the host institution, Instituto Gulbenkian de Ciência, and by the National Veterinary Authority (DGAV, Portugal; permit number 0421/000/000/2013). OXTR mutant zebrafish line (ZFIN ID: ZDB-ALT-190830-1) was generated and provided by Dr. Gil Levkowitz (Weizmann Institute of Science) using a TALEN-based genome editing system. The characterization of this line has been described in Nunes et al. (26).

All the experimental groups were formed at 4 days post-fertilization, based on the genotype of the progenitors, before they imprint for olfactory and visual kin recognition (27,28). To evaluate genotype-environment effects, fish were raised in groups according to the experimental design in **Figure 1A** and both female and males tested in adulthood (3 months old).

### Genotyping

At 3 months old, before behavioural screenings, genomic DNA was extracted from adult fin clips using the HotSHOT protocol (29). The genomic region of interest was amplified by PCR and sequenced to identify the focal fish in each group. The following primers were used: sense 5’-TGCGCGAGGAAAACTAGTT-3’, antisense 5’-AGCAGACACTCAGAATGGTCA-3’.

### Behavioural assays

#### Video acquisition

Fish were in a tank placed on top of an infrared lightbox and video-recorded either from above (shoal preference and social recognition tests) or laterally (group behaviour tests). Video acquisition was done with software Pinnacle Studio 14 (Corel Corporation, Ottawa, Canada). Shoal preference, social habituation and social recognition analyses were performed with EthoVision video tracking system (Noldus Information Technologies, Wageningen, The Netherlands) and group behaviour analyses were done with the open source FIJI image-processing package (30).

### Social preference and social habituation

The social preference test assesses the individual’s sociability by observing the interactions between conspecifics (22): a focal fish was placed in a central compartment (30×15×10 cm) of a three-compartment tank, separated by transparent and sealed partitions. A shoal of fish was placed in one of the lateral compartments (15×10×10 cm), while the other contained only water. To avoid any side bias the stimuli were balanced across trials. After an acclimatization period (10 min), the focal fish was released from a start box and allowed to explore the tank, while its behaviour was video-recorded for 10 min. The time spent by the focal fish near (less than two body lengths) each compartment was quantified and used to calculate the social preference score (*SP* = Time near shoal/ [Time near shoal + Time near empty]). A score above 0.5 indicates a preference for the shoal.

The social preference test was performed twice, with 24 hrs in between, and social preference scores of both tests were used to calculate the habituation index (*Hab. Score* = 1- [SP_Trial2_]/[SP_Trial1_ + SP_Trial2_]). A score above 0.5 reveal a decrease in preference to associate with conspecifics.

### Social recognition

The social recognition assay to evaluate short-term (i.e. 10 min retention) social memory was adapted from the procedure already developed in our lab for long-term (i.e. 24h retention) social memory in zebrafish (31), and has already been used successfully in a previous study (22). A focal fish was placed for 10 min in the central compartment of a three-compartment tank, separated by transparent and sealed partitions, to acclimatize. The focal fish was allowed to interact visually across partitions with 2 novel conspecifics for 10 min. After, both stimuli were removed, one was placed in the same compartment (familiar conspecific stimulus), while a novel conspecific was placed in the other compartment (novel conspecific stimulus). In a second 10 min interaction the time spent by the focal fish near each compartment (termed novel cue or familiar cue) was quantified and used to measure the preference for the novel (*Disc.Score*= Time near Novel/[Time near Novel + Time near Familiar]). A discrimination score of 0.5 indicates no preference between novel or familiar conspecifics.

### Shoaling behaviour

Shoaling behaviour is a common behaviour present in fish models and allows to determine complex interactions between individuals. Both focal fish and social partners were recorded in the home tanks (3.5L tank). Focal fish were tagged with fin clips for easy identification. The behaviours were video-recorded from side view for 10 min. Two components of shoaling behaviour were analysed in time bins of 8 seconds: (1) focal fish distance to the group centroid (social integration); and (2) the dispersion of the remaining shoal members as measured by their perimeter (social influence).

### Data analysis

Data were analysed using SPSS 25.0. All data sets were tested for departures from normality with Shapiro-Wilks test. Two factor univariate ANOVA were used for comparing multiple groups. All data sets were corrected for multiple comparisons. Tukey’s Test comparisons were used as post-hocs. Graphs were performed with GraphPad software.

### Ethical Approval

All experiments were performed in accordance with the relevant guidelines and regulations for the care and use of animals in research and approved by the competent Portuguese authority (Direcção Geral de Alimentação e Veterinária, permit 0421/000/000/2017).

## Acknowlegdements

We thank IGC Fish Facility for assistance on fish maintenance and Peter McGregor for comments and discussion on an earlier version of the manuscript. The authors declare no conflicts of interest related to this work. This work was funded by a Fundação para a Ciência e a Tecnologia (FCT) research grant (PTDC/BIA-ANM/0810/2014) and a FEDER grant (Lisboa-01-0145-FEDER-030627) awarded to RFO. ARN and MST were supported by Post-doc fellowships from Fundação para a Ciência e a Tecnologia (FCT, ARN: SFRH/BPD/93317/2013). Congento LISBOA-01-0145-FEDER-022170, co-financed by FCT (Portugal) and Lisboa2020, under the PORTUGAL2020 agreement (European Regional Development Fund). The authors declare no competing interests.

## Author contributions

R.F.O. and D.R. designed the research; D.R. and A.R.N. performed the research; D.R., A.R.N. and M.S.T. carried out the molecular lab work; A.S., J.B. and G.L. developed and validated the mutant zebrafish line; D.R. and M.S.T. carried out the statistical analyses; D.R. and R.F.O. wrote the paper and all authors critically revised the manuscript.

